# Grand paternal inheritance of an acquired metabolic trait induced by ancestral obesity is associated with sperm RNA

**DOI:** 10.1101/042101

**Authors:** Jennifer E Cropley, Sally A Eaton, Alastair Aiken, Paul E Young, Eleni Giannoulatou, Joshua WK Ho, Michael E Buckland, Simon P Keam, Gyorgy Hutvagner, David T Humphreys, Katherine G Langley, Darren C Henstridge, David IK Martin, Mark A Febbraio, Catherine M Suter

## Abstract

Parental exposure to an environmental challenge can induce phenotypes in offspring independent of the inherited DNA sequence. Whether such acquired traits can be inherited – i.e., can manifest in a generation beyond that exposed to the precipitating insult as germ cells – is unclear. Here we report a latent metabolic phenotype induced by paternal obesity that is inherited into a second generation, without germ cell exposure. Sons of obese male mice exhibit defects in glucose and lipid metabolism that are only unmasked by post-weaning dietary challenge, yet they transmit these defects to their own progeny (F2) in the absence of the challenge. F1 sperm exhibit changes in the abundance of several small RNA species, including diet responsive tRNA-derived fragments. These data suggest that induced metabolic phenotypes may be propagated for multiple generations through the actions of noncoding RNA.

## Introduction

Offspring phenotypes can be affected by parental health or parental environmental exposures, independent of variation in the inherited DNA sequence. Examples of such non-genetic transgenerational effects occur with a broad range of stressors, from dietary stress and toxin exposure to psychological stress or trauma (reviewed in ^1^). In some instances, induced phenotypes can be observed across multiple generations, but whether such observations represent true inheritance of an acquired trait, or merely a residuum of the original exposure, is not clear. This distinction has adaptive significance for organisms living in a changing environment, given the potential for epigenetic states to respond to environment and selection^2,3^.

In those cases in which induced phenotypes have been observed over more than one generation, determination of true inheritance is confounded by alternative scenarios. Persistence of an induced phenotype into grandoffspring of exposed mothers can generally be attributed to direct exposure of developing germ cells within the offspring. Such a scenario underpins several recent reports of programming by parental metabolism. For example, male mice exposed to a poor intrauterine environment can transmit metabolic defects to their own offspring^4^^−^^6^. An alternative scenario is serial programming of the induced phenotype over multiple generations: that is, when the phenotype induced in offspring mirrors the parental condition, and thus can in turn program defects in second generation offspring, *et cetera*. Serial programming resulting from a repeatedly compromised gestational environment may underlie most, if not all, examples of multigenerational maternal programming^7^.

It is currently unclear whether an induced metabolic phenotype can be transmitted into successive generations without the continued influence of the inducing stimulus; in other words, whether true, non-genetic transgenerational inheritance of a metabolic phenotype can occur via gametes that were never exposed^8^. True inheritance of the effects of parental exposures is arguably best studied through the male lineage, where confounding effects of the intrauterine environment can be excluded. It is known that perturbed paternal metabolism can induce metabolic defects in first-generation offspring^5,9−11^, but not whether the effects persist into second or subsequent generations. While epidemiological observations suggest this could be the case^12,13^, the heritability of programming is difficult to ascertain in human cohort studies, where genetic heterogeneity, lifestyle factors, and the necessarily retrospective nature of longitudinal studies are major confounders.

We have addressed the question of inheritance directly using an isogenic rodent model of obesity and pre-diabetes. We used a dominant genetic mouse model of obesity in which the obesogenic allele can be segregated away from the offspring. In this model, paternal obesity induces a latent metabolic phenotype in first-generation sons that is unmasked by exposure to a high fat Western style diet. By breeding sons maintained on a healthy diet (exposed to paternal obesity but metabolically normal), we find that the latent phenotype induced by paternal obesity is inherited into a second, unexposed, generation. The inheritance is associated with changes to the small RNA profile of sperm in first-generation offspring, despite the sperm developing in a normal metabolic environment.

## Results

### Sons of obese sires carry a latent predisposition to hepatic insulin resistance

To model paternal obesity, we used C57BL/6 mice carrying the dominant *agouti viable yellow* (*A^vy^*) mutation, which produces a phenotype of yellow fur, hyperphagia, and maturity-onset obesity^14^. Obese yellow *A^vy^*/*a* males are hyperinsulinemic but not hyperglycaemic (Supplementary Fig. 1), modelling the majority of obese men of reproductive age who are insulin resistant but not frankly diabetic^15^. Obese *A^vy^*/*a* heterozygote sires can be mated to congenic lean *a*/*a* dams to generate *a*/*a* offspring that are genetically identical to offspring from *a*/*a* matings. In this way, we generated two isogenic groups of F1 offspring differing only in the *A^vy^* genotype, and the phenotypes, of their sires (see Supplementary Fig. 1 for breeding strategy and Supplementary Table 1 for breeding statistics). Offspring of each lineage were randomly assigned at weaning to either control diet (CD) or to a Western-style diet (WD) high in saturated fat and sugar (Supplementary Table 2).

When maintained on either CD or WD, weight trajectories of male *a*/*a* offspring of obese *A^vy^*/*a* sires (PatObF1) were no different to offspring of lean sires (Fig. 1a). But while all animals gained weight on WD, PatObF1 animals gained more gonadal fat than controls (Fig. 1b), suggesting that paternal obesity confers an increased sensitivity to the adipogenic effects of a high-fat diet. Similarly, glucose tolerance was normal in all mice when maintained on CD (Fig. 1c), but after only three weeks on WD, PatObF1 males show a significantly impaired response in an intraperitoneal glucose tolerance test (Fig. 1d). The same degree of intolerance was observed whether the offspring were generated from an overnight timed mating or continuous co-housing of sire and dam (Fig. 1d); this indicates that the programmed defect was transmitted at the time of mating, rendering programming via social effects of the obese sire on the dam or offspring improbable. PatObF1 males also had exacerbated hyperinsulinemia in response to WD (Fig. 1e), together with significantly elevated intrahepatic triacylglycerol (TAG), and the insulin resistance driver diacylglycerol (DAG) (Fig. 1f, g)^16^. These data indicate that paternal obesity programmed a latent predisposition to hepatic insulin resistance that was unmasked by challenge with a Western diet. We observed this programming only in males: for the duration of the experiment, female PatObF1 exhibited normal glucose homeostasis on both CD and WD (Supplementary Fig. 2).

**Fig. 1.**
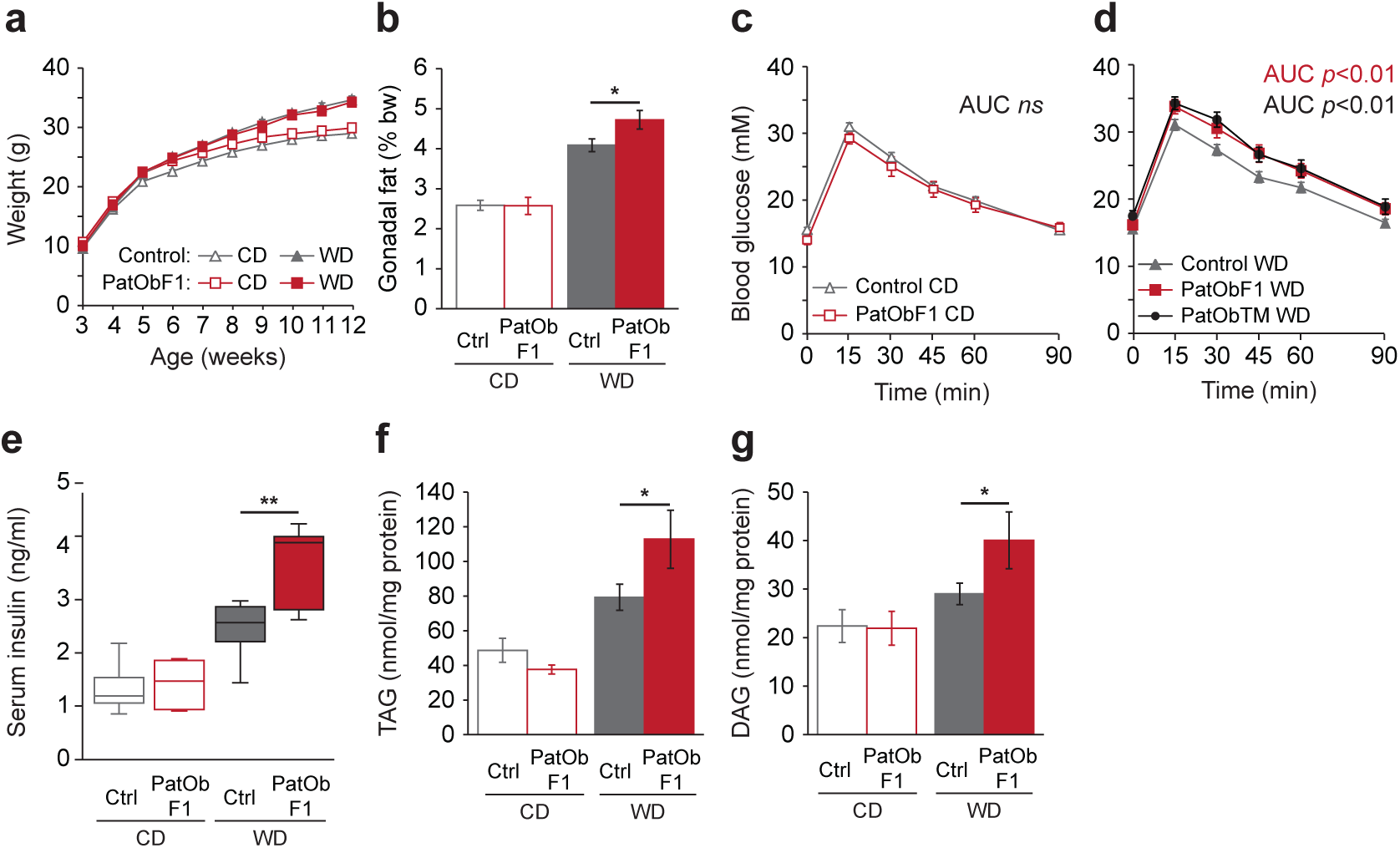
Paternal obesity programs latent metabolic defects in F1 male offspring. (**a**, **b**) Body weights (**a**) and relative gonadal fat weights at 12 weeks (**b**) of Control and PatObF1 offspring fed control diet (CD) or Western diet (WD); CD: Control *n* = 46, PatOb *n* = 16; WD: Control *n* = 43, PatOb *n* = 19. (**c**, **d**) Glucose tolerance tests on six-week old CD offspring (**c**) and WD offspring (**d**), including those from time-mated obese sires (PatObTM, *n* = 16; other animal numbers as in **a**). (**e**) Serum insulin in 12-week old offspring; CD: Control *n* = 9, PatOb *n* = 5; WD: Control *n* = 8, PatOb *n* = 5. (**f**, **g**) Hepatic triacylglyceride (TAG, **f**) and diacylglyceride (DAG, **g**) levels in 12-week old offspring; CD: Control *n* = 7, PatOb *n* = 8; WD: Control *n* = 8, PatOb *n* = 8. Error bars represent SEM; * *p* < 0.05, ** *p* < 0.01.

### The latent metabolic phenotype induced by paternal obesity is inherited from F1 sons into F2 grandsons

We then asked whether the defects programmed by paternal obesity could be inherited, that is, passed into a second, unexposed generation. Here the latency of the phenotype in F1 provides a material advantage in distinguishing inheritance from serial programming, since the germ cells and sperm of F1 mice on CD were not exposed to metabolic dysfunction. We found that the latent metabolic phenotype was inherited through the male line into F2: the PatObF2 males faithfully recapitulated the phenotype of their F1 sires. Again the weight trajectories of PatObF2 grandsons were no different from control (Fig. 2a), but PatObF2 had a greater gonadal fat mass on WD (Fig. 2b). With the WD challenge, PatObF2 grandsons exhibited the same pathognomonic signs of hepatic insulin resistance displayed by PatObF1 sons: marked glucose intolerance (Fig. 2d), hyperinsulinemia (Fig. 2e) and elevated hepatic TAG and DAG (Fig. 2f, g). The only contrast with the phenotype of F1 is that PatObF2 exhibited hyperinsulinema and a mild glucose handling impairment even on CD (Fig. 2 c, e), suggesting that the phenotype inherited to F2 is slightly more overt than in F1. Taken together, these data indicate that the latent predisposition to metabolic defects induced by paternal obesity is inherited into a second, unexposed, generation. The germ cells that gave rise to the F2 generation developed while the F1 mice were gestating in a normal intrauterine environment, and matured inside an F1 individual that was not obese and had no overt metabolic disturbance, arguing against serial programming as the mechanism of transmission. Thus we have observed true, non-genetic transgenerational inheritance of an acquired metabolic phenotype induced by ancestral obesity.

**Fig. 2.**
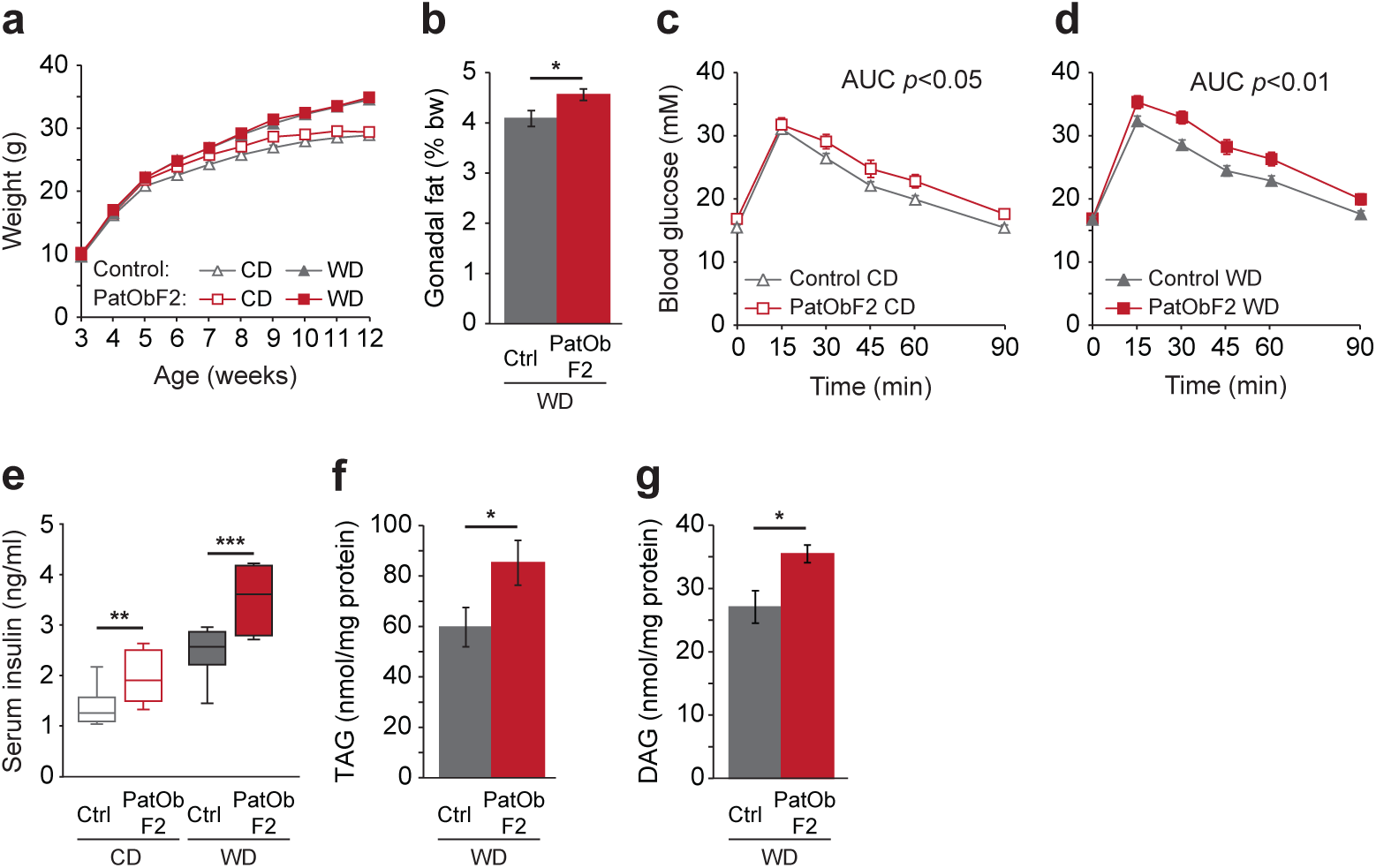
Metabolic programming by paternal obesity is inherited by a second generation. (**a**) Body weights of Control and PatObF2 offspring fed control diet (CD) or Western diet (WD); CD: Control *n* = 46, PatObF2 *n* = 17; WD: Control *n* = 43, PatObF2 *n* = 17. (**b**) Relative gonadal fat weights in 12-week old mice; Control and PatObF2 animal numbers as in **a**. (**c**, **d**) Glucose tolerance tests on six week old CD (**c**) and WD (**d**) mice; animal numbers as in **a**. (**e**) Serum insulin in 12-week old mice; CD: Control *n* = 9, PatObF2 *n* = 8, WD: Control *n* = 8, PatObF2 *n* = 7. (**f**, **g**) Liver triacylglyceride (TAG, **f**) and diacylglyceride (DAG, **g**) levels in 12-week old mice on WD; Control, *n* = 8, PatObF2 *n* = 7. Error bars represent SEM; * *p* < 0.05, ** *p* < 0.01, *** *p* < 0.001.

### Inheritance of the latent metabolic phenotype into F2 is associated with changes in F1 sperm RNA content

The characteristics of the inheritance we observe implicate an epigenetic inheritance system. Small noncoding RNAs direct epigenetic states in the germline^17,18^ and have been implicated in invertebrate models of epigenetic inheritance^19^^−^^22^. Recently two separate studies have shown that the small RNA content of murine sperm is sensitive to dietary perturbations: both a low-protein diet and a high-fat diet are linked to changes in the abundance of specific small RNA species in exposed sperm^11,23^. We asked whether small RNA perturbations may also underlie the *inheritance* of the induced metabolic phenotype we observe. To do this we examined the small RNA content of sperm from F1 offspring of obese sires fed the control diet: these males are able to transmit the latent phenotype, but do not exhibit it. We compared the sperm small RNA profiles with those from control-fed control males.

The distribution of small RNA lengths and biotypes between the two groups was very similar (Fig. 3a,b), but within two biotypes – miRNA and tRNA-derived fragments (tRFs) – there were major differences in abundance of some species. Using stringent thresholds (only considering miRNA with counts >200 per million, and adjusted *p*-value of < 0.05) we found 24 miRNAs with altered expression in PatObF1 (Figure 3c, Table 1). One sperm miRNA, miR-10, exhibited dramatic changes, increasing 2.5-fold to constitute ∼25% of all sperm miRNA in PatObF1 (Fig. 3). Both miR-10 isotypes, miR-10a and miR-10b (encoded by different genes) were significantly elevated. Differing only by one nucleotide which is outside of the seed sequence, these two miRs target the same set of mRNAs (Supplementary Table 3); these targets are heavily and significantly enriched for functions related to the regulation of transcription (Fig 3e).

Around a quarter of all sperm small RNAs were tRFs of two predominant species, tRF5-GluCTC and tRF5-GlyGCC: collectively, sequences of these two isotypes contributed ∼90% of all sperm tRFs. The relative ratios of these tRFs were inverted in PatObF1 sperm relative to control sperm, with an ∼30% reduction in tRF5-GluCTC and a corresponding increase in tRF5-GlyGCC (Fig. 3f). Another two tRFs, ValCAC and HisGTG were also significantly elevated in PatObF1 sperm, although these tRFs were much less abundant, constituting less than 5% of all sperm tRFs. Both tRF5-GluCTC and-GlyGCC display precise cleavage at the 5’ side of the anticodon loop of the mature tRNA to generate fragments of predominantly 30 and 31 nucleotides, respectively (Fig. 3g). GluCTC and GlyGCC are the same tRFs that were reported as perturbed in sperm by exposure to an altered diet^11,23^.

**Fig. 3.**
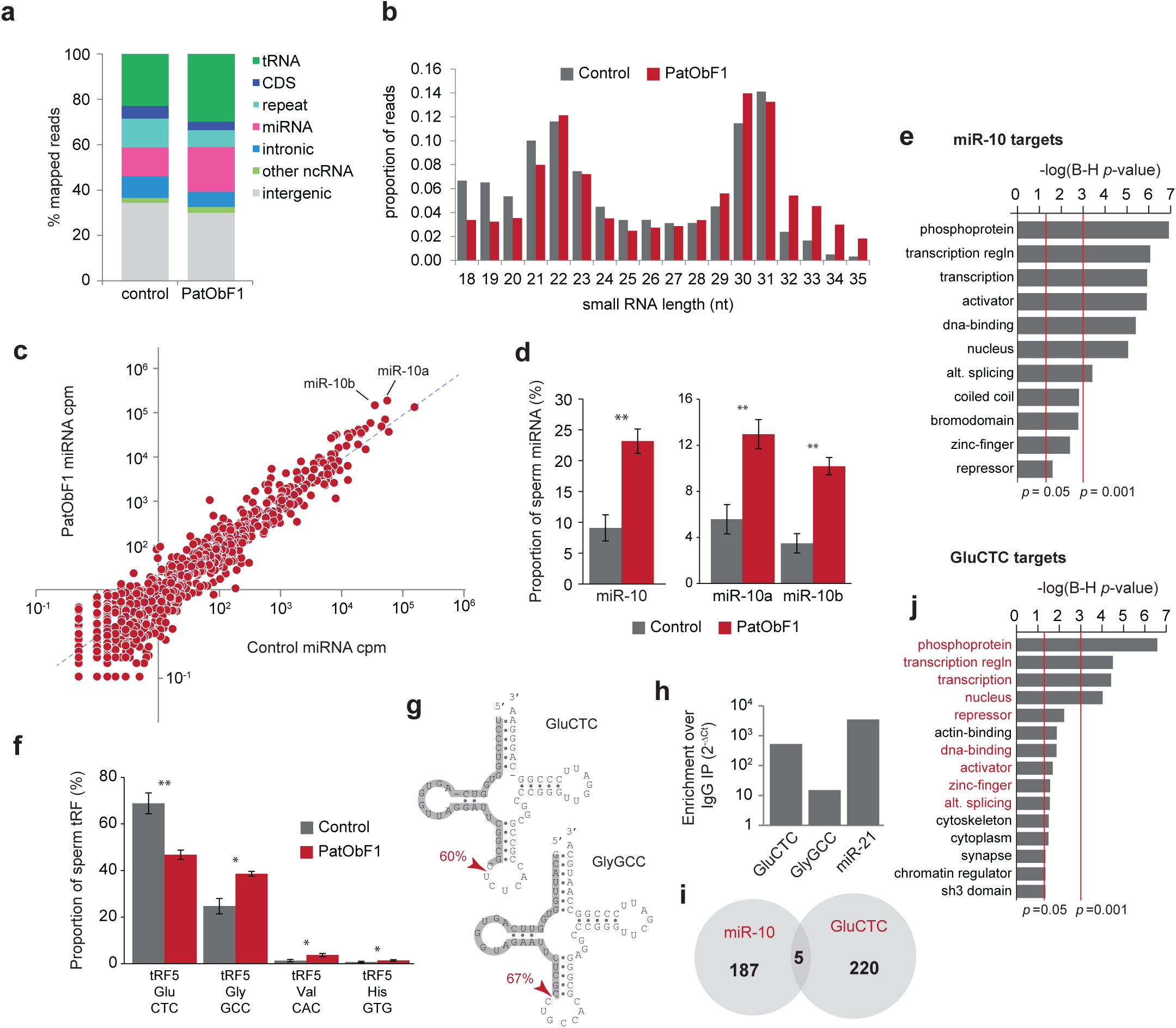
Paternal obesity alters small RNAs in the sperm of F1 sons. Small RNA biotype (**a**) and size distribution **(b)** of all 18-35nt sRNA reads in Control and PatObF1 sperm. **(c)** Scatterplot of sperm miRNA abundance from Control (x-axis) versus PatObF1 (y-axis). **(d)** miR-10 as a fraction of all sperm miRNA in PatObF1 (red) relative to Control (grey); panel at right shows levels of the two miR-10 isoforms ** *q* < 0.01 (**e**) Gene ontologies significantly enriched in the targets of miR-10. **(f)** tRFs significantly altered in PatObF1 sperm (red) relative to control sperm (grey) representated as proportion of all sperm tRFs * *p* < 0.05, ** *p* < 0.01 (**g**) Clover leaf structures of GluCTC (top) and GlyGCC (bottom); dominant sperm tRF isoform shaded grey. (**h**) Results of Taqman RT-PCR on RNA purified from an Ago2 immunoprecipitation (IP) showing tRF enrichment in the IP over the negative control IgG IP; miR-21 is a positive control.**(i)** Individual gene targets of miR-10 and tRF5-Glu-CTC show little overlap. (**j**) Gene ontologies significantly enriched in the targets of tRF5-Glu-CTC; ontologies in common with miR-10 are shown in red.

**Table 1:** microRNAs with altered abundance in sperm of PatObF1^a^

| microRNA | Average counts per million<sup>b</sup> |  | Fold Change | q-value |
| --- | --- | --- | --- | --- |
|  | Control | PatOb |  |  |
| mmu-miR-204 | 717 | 3,749 | **5.23** | 0.0002 |
| mmu-miR-182 | 694 | 2,298 | **3.31** | 0.0043 |
| mmu-miR-429-3p | 386 | 1,182 | **3.06** | 0.0003 |
| mmu-miR-10b | 35,008 | 101,932 | **2.91** | 0.0012 |
| mmu-miR-146b | 566 | 1,634 | **2.88** | 0.0192 |
| mmu-miR-200a | 280 | 793 | **2.83** | 0.0004 |
| mmu-miR-148a-3p | 7,619 | 21,046 | **2.76** | 0.0033 |
| mmu-miR-200a-3p | 6,752 | 18,443 | **2.73** | 0.0033 |
| mmu-miR-200b-3p | 8,257 | 22,186 | **2.69** | 0.0039 |
| mmu-miR-451a | 632 | 1,666 | **2.64** | 0.0257 |
| mmu-miR-183 | 1,226 | 2,985 | **2.44** | 0.0088 |
| mmu-miR-10a | 55,867 | 129,719 | **2.32** | 0.0088 |
| mmu-let-7e | 628 | 1,205 | **1.92** | 0.0267 |
| mmu-miR-210-3p | 350 | 648 | **1.85** | 0.0266 |
| mmu-miR-187-3p | 245 | 378 | **1.54** | 0.0438 |
| mmu-miR-328-3p | 1,661 | 582 | **0.35** | 0.0265 |
| mmu-miR-676-3p | 2,003 | 699 | **0.35** | 0.0262 |
| mmu-miR-378a-3p | 5,059 | 1,740 | **0.34** | 0.0246 |
| mmu-miR-669c | 903 | 309 | **0.34** | 0.0433 |
| mmu-miR-92a-3p | 12,593 | 4,390 | **0.35** | 0.0341 |
| mmu-miR-425 | 5,340 | 1,592 | **0.30** | 0.0405 |
| mmu-miR-127-3p | 1,470 | 360 | **0.24** | 0.0033 |
| mmu-miR-184-3p | 7,487 | 1,808 | **0.24** | 0.0103 |
| mmu-miR-34b-3p | 12,678 | 2,073 | **0.16** | 0.0052 |

<sup>a</sup> Where sequencing abundance > 200 reads per million and $q < 0.05$ ; <sup>b</sup> Reads per million mapped miRNA reads.

Mature sperm are transcriptionally and translationally inert, so any function of the sperm-borne tRFs and miRNAs is likely to manifest post-fertilisation. Recently reported diet-responsive tRFs induce transcriptional changes in the early embryo; whether this is a direct or indirect transcriptional effect is unclear. Available evidence indicates that tRFs reside almost exclusively in the cytoplasm^24^ where they can associate with polyribosomes and affect translation^25^^−^^27^. The majority of miRNAs are also found in the cytoplasm and functionally associate with Ago2. Ago2 is highly expressed in the oocyte and early embryo and is required for the maternal-to-zygotic transition (MZT)^28^. We asked whether the two sperm tRFs can also associate with Ago2, and found that tRF5-GluCTC, and to a lesser extent tRF5-GlyGCC, was enriched in an Ago2 pulldown (Fig. 3h). This indicates that at least tRF5-GluCTC binds Ago2, signifying its potential to act in a miRNA-like manner in the zygote. Using a miRNA ‘seed’ sequence-based algorithm we identified potential tRF5-GluCTC targets: the individual predicted targets of tRF5-GluCTC shared very little overlap with those of miR-10, yet remarkably resided in strikingly similar transcriptional regulation ontologies (Fig. 3i,j, Supplementary Table 3). Using publicly available datasets of murine early embryonic transcription^29^ we found that 114/220 miR-10 targets and 77/191 tRF-5-GluCTC targets are expressed in the oocyte; approximately half of these targets are reduced by at least two-fold at the 4-cell stage relative to the oocyte, consistent with a repressive post-transcriptional function of miR-10 and tRF-5-GluCTC.

## Discussion

We have identified an acquired metabolic phenotype induced by paternal obesity that persists for two generations, independent of the inherited genotype, the gestating environment, and continued exposure to the inducing stimulus. The characteristics of our model – in particular the latency of the induced phenotype – strongly suggest that the transmission we observe represents true inheritance of an acquired phenotype. While there has been one previous report of a paternally-induced and inherited behavioural phenotype in mice^30^, to our knowledge, this is the first clear demonstration of the inheritance of a paternally-induced metabolic phenotype in vertebrates. Our findings have immediate relevance to the evolution of human disease and suggest a mechanism by which the memory of ancestral exposures may be maintained through multiple generations.

In our model, obese founder males had a syndrome of obesity and hyperinsulinemia, but the F1 males that gave rise to the F2 generation did not. When maintained on a regular chow diet, F1 males were effectively indistinguishable from isogenic controls in terms of weight, glucose metabolism, insulin levels, and hepatic lipids; this argues strongly against the serial transmission of the metabolic defects from F1 to F2 and implies epigenetic inheritance. The possibility that the latent phenotype was transmitted by behavioural factors is also unlikely: dams partnered with obese sires were no heavier than dams of control sires, indicating that they did not acquire the hyperphagia of their partners, and the programmed phenotype was just as robust when the influence of the sire was restricted to an overnight timed-mating. The most parsimonious explanation for the transmission of the latent metabolic phenotype is transgenerational epigenetic inheritance: some mark or memory of paternal obesity passaged via the male germline, implicating sperm in the transmission.

We found alterations to multiple prominent small RNA species in the sperm of mice born of obese fathers. The idea that sperm RNA is involved in transgenerational epigenetic inheritance is well established in invertebrate systems. In the fly, small RNAs influence the transgenerational inheritance of maternal immunity to paternally-introduced transposons^22^. In the nematode *C. elegans*, small noncoding RNAs are implicated in multiple models of transgenerational inheritance^19,20,31^ including a metabolic starvation-induced response^21^. There is more limited, and somewhat controversial, evidence for small RNA involvement in inheritance of vertebrate phenotypes: injection of sperm RNA into fertilised oocytes has been reported to recapitulate several induced phenotypes, including white tail tipping^32^, cardiac hypertrophy^33^ and fear conditioning^34^.

Molecular changes to mouse sperm (including alterations to DNA methylation, miRNAs, and tRFs), have been described in response to environmental factors, including dietary perturbations ^4,6,10,11,23,34^, but in these studies sperm were directly exposed to the initiating stimulus, either as mature gametes or as developing germ cells. In this study we did not examine the directly exposed sperm (i.e. that from the obese founder males), because identification of any agent of inheritance may have been confounded by changes induced by direct exposure of the sperm to the syndrome of obesity and pre-diabetes. It is remarkable that we find alterations to the same tRF species as have been recently reported to be male diet-sensitive^11,23^. The sperm from F1 males in our study were not exposed to obesity or altered diet, or indeed to any apparent insult at all, but they still transmitted the induced phenotype to the F2 generation. Together these data provide compelling evidence implicating tRFs and other small RNAs in the inheritance of acquired metabolic traits in vertebrates.

Given that mature sperm are transcriptionally and translationally inert, it is likely that sperm small RNAs function post-fertilisation. Recent work in *C. elegans* found that around half of all small noncoding RNAs in the one-cell embryo are paternally derived^35^. This raises the possibility that sperm-borne small RNAs play a role in very early development prior to, or during, the maternal-to-zygotic transition^36^. Both of the recent studies that identified diet-regulated sperm tRFs found changes in early embryonic transcriptional invoked a direct transcriptional interference mechanism for sperm-borne tRFs^11,25^. However, available evidence indicates that tRFs reside almost exclusively in the cytoplasm^24^ where they can associate with polyribosomes and affect translation^25^^−^^27^. Our data suggest a post-transcriptional regulatory function that indirectly affects transcription. The targets of the two small RNA species in sperm most affected by paternal obesity, tRF5-GluCTC and miR-10, are heavily enriched in transcriptional regulatory functions, implying that changes in their abundance in sperm may perturb transcriptional regulation in the developing embryo post-transcriptionally. These targets are significantly downregulated by the 4-cell stage, when the MZT is complete; these early perturbations may set in train an altered physiology and reduced resilience to environmental insults (such as over-nutrition) in later life.

If the phenomenon we observe here translates to humans, our findings suggest that persons with obese paternal ancestry could avoid the deleterious effects of metabolic programming with dietary intervention, yet still propagate the propensity for metabolic dysfunction to their offspring. The implications for public health are obvious. Given that a wide range of phenotypes can be induced by a variety of paternal factors^37^, it is not unreasonable to suppose that such epigenetic inheritance may propagate the risk of many human diseases. How the associated RNA perturbations induced by paternal obesity are maintained in germ cells and gametes an entire life cycle beyond the inducing stressor is currently a mystery.

## Materials and methods

### Mice and diets

All animals were handled in accordance with good practice as defined by the National Health and Medical Research Council (Australia) Statement on Animal Experimentation, and requirements of state government legislation. The study was approved by the Garvan/St Vincent’s Animal Ethics Committee (12/09 and 13/35).

The *A^vy^* mice used in this study were descended from an isogenic C57BL/6 colony at Oak Ridge National Laboratories (Oak Ridge, TN, USA), and have been maintained at the Victor Chang Cardiac Research Institute since 2001. *A^vy^*/*a* mice display a variety of phenotypes from obese yellow through degrees of coat-colour mottling to lean agouti^38^. To avoid any effects of ancestral obesity in our stock *A^vy^* colony, we maintain it by breeding only lean agouti *A^vy^*/*a* mice to congenic *a*/*a* mice. The breeding strategy is outlined in Supplementary Fig. 1, and breeding statistics in Supplementary Table 1. For experimental mice, yellow *A^vy^*/*a* males were selected from the stock colony for mating to *a*/*a* females. For control mice, we paired *a*/*a* males and females from the stock colony and selected their offspring as founders, so that control founders were at least two generations removed from any heritable metabolic effects of the *A^vy^* allele. Control mice were generated contemporaneously with PatOb mice over three generations, and pooled for analysis, after we ensured that metabolic characteristics of different generations of control mice were equivalent.

The mice in this study were produced over the course of two years in the same facility. To maximise breeding output (obese yellow *A^vy^*/*a* mice are poor breeders), in general males were allowed to cohabit with females throughout the breeding period. To control for social effects on the females, some animals were time-mated: males were placed in the female’s cage overnight and removed early the next morning along with any faeces; this was repeated up to three times until a positive plug was observed.

All breeding mice were fed *ad libitum* on NIH-31 control diet from weaning (5% w/w fat, 13.5 MJ/kg). *a*/*a* experimental offspring were randomly assigned at weaning to either SF00-219 high-fat diet designed to mimic a Western fast-food diet (WD, equivalent to Harlan Teklad TD88137; 22% w/w fat (40% digestible energy), 19.4 MJ/kg, see Supplementary Table 2) or matched normal fat control (CD; 6% w/w fat, 16.1 MJ/kg, see Supplementary Table 2). Any *A^vy^*/*a* littermates in F1 were culled at weaning. Feeds were manufactured by Specialty Feeds (Glen Forrest, WA, Australia).

Experimental mice were culled at 12 weeks of age; metabolic organs were harvested and weighed and blood collected, and sperm were harvested from some animals (see below).

### Metabolic profiling

Glucose tolerance testing was performed on six week old mice; the tester was unaware of the group allocation of the individuals being tested. Mice were fasted for 6 hours and given an intraperitoneal injection of 1.5 g glucose/kg weight. Blood glucose levels were measured at 0, 15, 30, 45, 60 and 90 minutes using an AlphaTrak glucose monitor (Abbott). Animals were excluded from the analysis if their blood glucose did not increase by at least 75% by 15 minutes post-injection, or if excessive fighting between animals before injection raised the basal glucose measurement more than two standard deviations above the group mean.

Serum insulin levels were measured in 12 week blood samples using the Mouse Insulin ELISA kit (Millipore); the tester was unaware of the group allocation of the samples being tested.

Lipidomic profiling was performed on 12 week liver samples, as described previously; the tester was blinded to the group allocation of samples. Samples more than two standard deviations from the group mean were excluded.

### Sperm isolation and RNA preparation

Mature motile sperm were isolated from the cauda epididymis of 12 week old experimental mice on CD. Caudae were dissected and sliced open, and sperm allowed to swim out into DMEM media on a heating pad. Media was collected after 30 minutes and either snap frozen in liquid nitrogen or processed immediately. After thawing, samples were centrifuged at 50 *g* to remove any tissue debris. The supernatant was collected and sperm were centrifuged for 3 minutes at 2500 *g*, washed twice in PBS, incubated in somatic cell lysis buffer (0.5% Triton-X, 0.1% SDS) for five minutes, washed again and resuspended in PBS. A proportion of the sperm were examined microscopically for the presence of somatic cells: only sperm samples with no somatic cells visible in 500-1000 sperm visualised were used for RNA analysis. RNA was extracted from pelleted sperm using Trizol; sperm lysis was assisted by the addition of 200mM β-mercaptoethanol, and repeated passage through a 29g needle.

### Small RNA-Seq

Barcoded small RNA sequencing libraries were prepared from ∼100 ng sperm total RNA using the NEBNext Small RNA for Illumina kit (NEB). Ligation of the 3’ adapter was allowed to proceed overnight to encourage maximal ligation of 3’O-methylated RNAs. Libraries were sequenced on an Illumina HiSeq 2500 in Rapid Run mode.

Fastq files were trimmed using Cutadapt (v1.6) to lengths of 18-51nt, with 10% adapter sequence error allowed. Trimmed reads were aligned to genome build mm 10 using Bowtie (v1.1.1) with parameters:-sam-n 1-l 18-e 70 (one mismatch allowed in an 18 base seed and a maximum sum of quality scores for mismatched bases of 70). Multimappers were randomly assigned among best hits. Mapped reads were annotated against miRNAs (mm10 mirbase20; mirbase.org), tRNAs (UCSC golden path), nested repeats (UCSC golden path), refgene (UCSC golden path) using custom perl scripts; annotation ties were broken by considering (in order) the orientation of the read relative to the genetic element, biotype hierarchy (according to the list above) and the number of bases overlapping the genetic element. For miRNA comparisons we used generalised linear modelling with EdgeR^40^, applying an abundance threshold of 0.02% total miRNA (i.e. 200 reads per million mapped miRNA reads).

### FLAG-Ago2 immunoprecipitation

HeLa cells were transfected with a FLAG-Ago2 construct in p3XFLAG-CMV-10 plasmid (Sigma). After 48 hours, ∼5x10^6^ cells were lysed and the lysate incubated with Anti-Flag M2 antibody or isotype control IgG (Sigma) bound to Dynabeads Protein G (Life Technologies). After washing, 10% of the immunoprecipitation was removed for Western blotting for Flag-Ago2, and the remaining 90% was used for small RNA quantification. For RNA preparation, beads were treated with Proteinase K and removed, and RNA was prepared using Trizol.

tRF5-GluCTC, tRF5-GlyGCC and hsa-miR-21-5p were assayed in HeLa cell total RNA (to confirm tRF expression) and IP RNA (to test for Ago2 enrichment) by Taqman MicroRNA assay (Life Technologies).

### miRNA/tRF target and GO analysis

To generate lists of target genes for miR-10a/b and tRF5-GluCTC, we used TargetScan v4.0, which allows input of a user-defined seed sequence^41^. We used DAVID ^42,43^ to determine significantly enriched ontologies. For analysis of miRNA/tRF target expression in oocytes and early embryo, we used data from DBTMEE^29^ using an abundance threshold of 3 FKPM.

### Statistical analyses

Weights trajectories were fitted using a linear mixed model^44^ and comparisons performed with a Wald test. For metabolic measurements (GTT, serum insulin, liver lipids), groupwise comparisons were performed with one-way ANOVA. Because sample size was not equivalent between groups, we confirmed the homogeneity of variances using Levene's test. Where variances were unequal, Welch's modified F statistic was used to assess significance; equivalent conclusions were obtained using the Welch ANOVA and the standard ANOVA. Post-hoc pairwise comparisons between groups of interest were performed with a one-sided Student’s t-test.

## Acknowledgements

Raw sequencing data has been deposited in the NCBI Sequencing Read Archive under study accession SRP051542. J.E.C. is supported by an Australian Research Council (ARC) DECRA Fellowship (DE120100723). G.H. is supported by an ARC Future Fellowship (FT110100455). M.A.F. is supported by an National Health and Medical Research Council (NHMRC) Senior Principal Research Fellowship (APP1021168). C.M.S. is supported by an ARC Future Fellowship (FT120100097). The study was supported by the Victor Chang Cardiac Research Institute, and in part by ARC DP120100825 and ARC DP130103027.

## Author contributions

C.M.S. and J.E.C. conceived and designed the project with input from S.A.E. and D.I.K.M.; S.A.E. and A.A. performed animal husbandry and metabolic analyses, with assistance from P.E.Y., J.E.C. and M.E.B.; J.E.C. performed small RNA profiling; S.P.K. performed Ago2 immunoprecipitation and Taqman assays; P.E.Y., D.T.H, E.G. and J.W.K.H performed bioinformatics analysis; K.G.L. and D.C.H. performed mass spectrometry; C.M.S., J.E.C., M.F., G.H. and D.I.K.M. analysed and interpreted data; C.M.S., J.E.C. and D.I.K.M. wrote the paper, with input from all authors.

## Competing financial interests

The authors declare no competing financial interests.

**Supplementary Figure 1.**
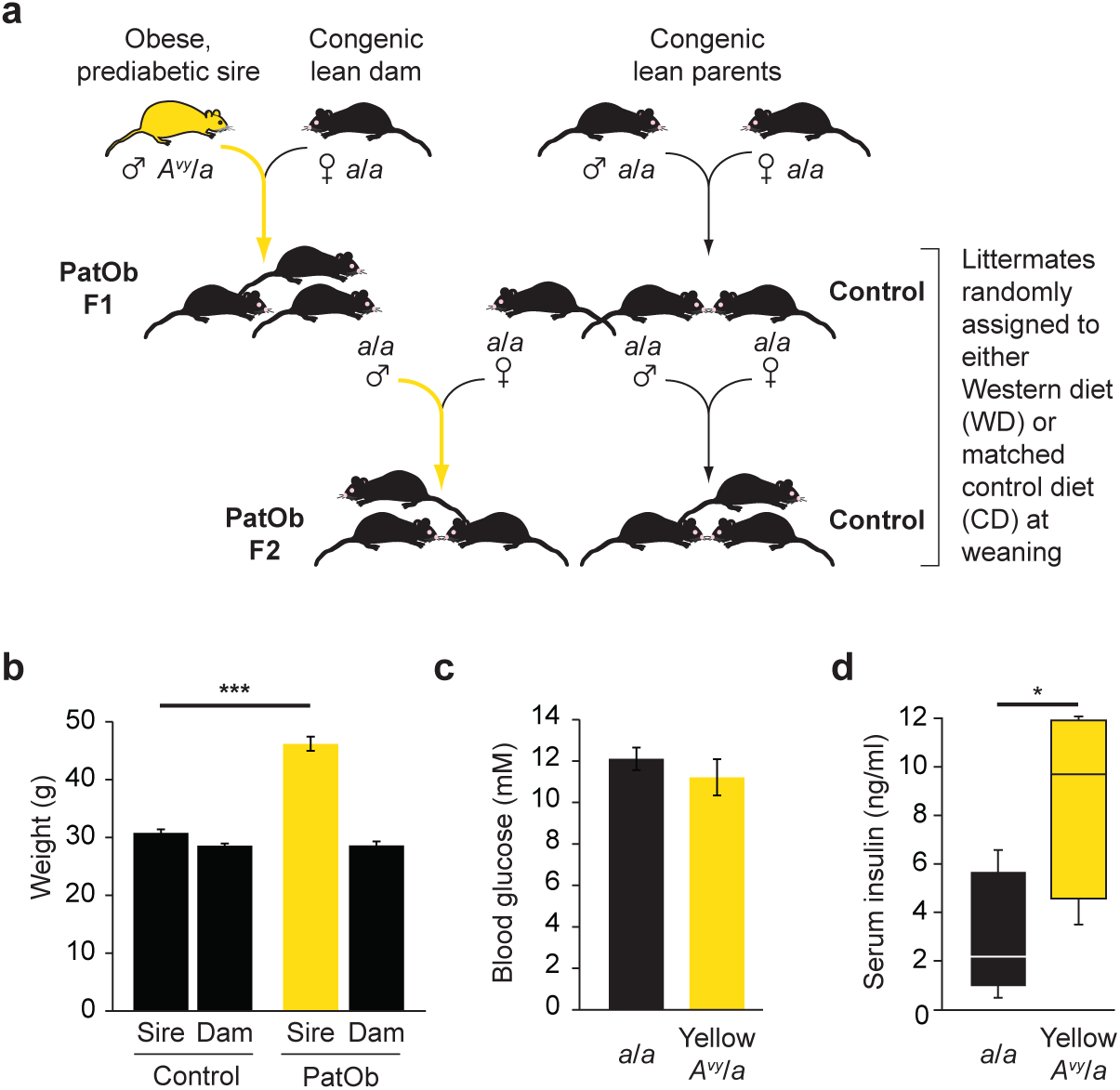
Experimental design and metabolic characteristics of obese founder males. (**a**) Schematic diagram showing breeding strategy. Obese yellow *A^vy^*/*a* males were mated with congenic lean *a*/*a* dams to generate PatObF1. PatObF1 males were mated with Control F1 daughters, generated by mating lean *a*/*a* mice, to generate PatObF2. For simplicity, *A^vy^*/*a* offspring of obese yellow *A^vy^*/*a* sires are not shown. (**b**) Weights of lean *a*/*a* sires and their *a*/*a* dams (Control, *n* = 62) and of obese yellow *A^vy^*/*a* sires and their *a*/*a* dams (PatOb, *n* = 21) measured one week after birth of offspring. (**c**) Blood glucose of lean *a*/*a* and obese yellow *A^vy^*/a males at 12 weeks of age (*n* = 12). (**d**) Serum insulin of lean *a*/*a* and obese yellow *A^vy^*/*a* males at 12 weeks of age (*n* = 5). Error bars represent SEM; * *p* < 0.05, *** *p* < 0.001.

**Supplementary Figure 2.**
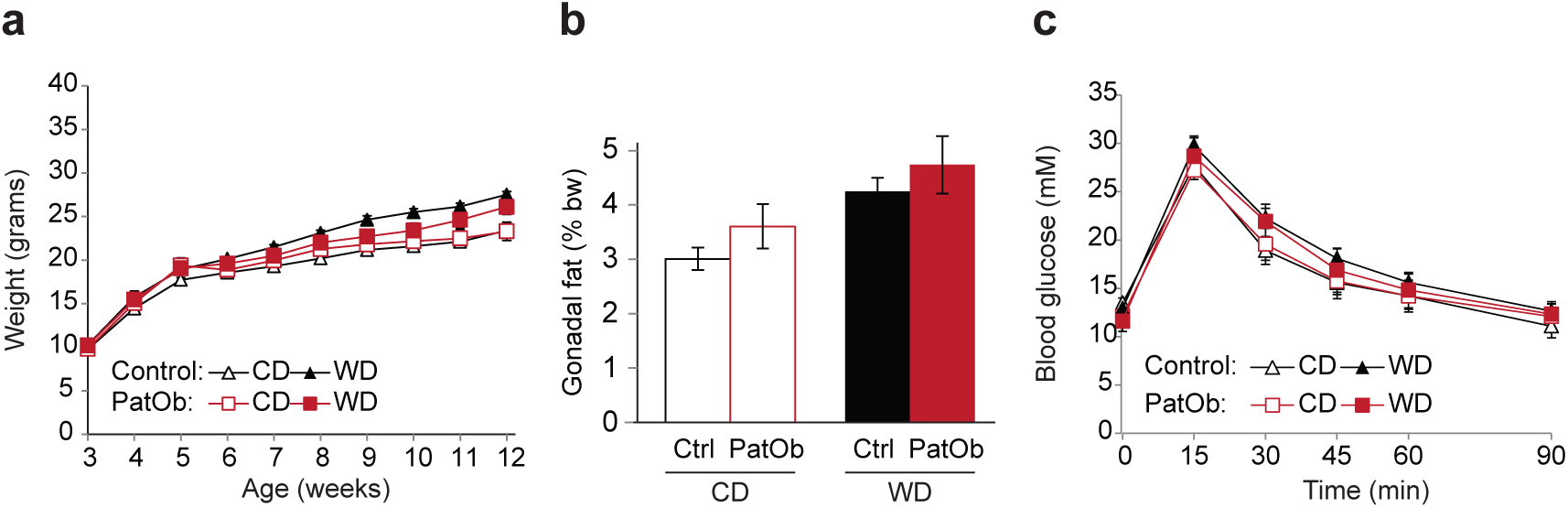
Female F1 offspring of obese sires are not fatter or predisposed to glucose intolerance. (**a**) Post-weaning weight trajectories of female offspring of obese sires (PatOb) or lean sires (Control), fed either Control diet (CD) or a Western diet (WD) (Control, *n* = 19–20; PatOb, *n* = 7). (**b**) Gonadal fat weight for female offspring as in **a**. (**c**) Glucose tolerance test at six weeks of age for F1 females, (Control, *n*=16–19; PatOb, *n* = 7).

**Supplementary Table 1:** Breeding statistics.

| Offspring Group | Sire genotype | Dam genotype | Number of sires | Number of litters | Average litter size<br>± SEM | Average sire age at birth<br>± SEM (d) | Average sire weight<br>± SEM (g)<sup>a</sup> |
| --- | --- | --- | --- | --- | --- | --- | --- |
| Control | *a/a* | *a/a* | 15 | 23 | 6.8 ± 0.4 | 123 ± 9 | 31.1 ± 0.9 |
| PatObF1 | *A<sup>vy</sup>/a* | *a/a* | 8 | 12 | 7.4 ± 1.1 | 126 ± 11 | 48.6 ± 1.2*** |
| PatObF2 | *a/a* | *a/a* | 6 | 15 | 6.9 ± 0.6 | 121 ± 12 | 33.5 ± 1.3 |

<sup>a</sup> sire weight at 7dpp
\*\*\* $p < 0.0001$ v Control

**Supplementary Table 2:** Diets used in this study.

|  | **Western**<br>**diet**<br>**(SF00-219)** | **Control**<br>**diet**<br>**(SF04-057)** |
| --- | --- | --- |
| Protein | 19.0% | 19.0% |
| Total Fat | 21.0% | 6.0% |
| Crude Fibre | 4.7% | 4.7% |
| AD Fibre | 4.7% | 4.7% |
| Digestible Energy | 19.4 MJ/Kg | 16.1 MJ/Kg |
| % Total calculated<br>digestible energy from<br>lipids | 40.0% | 14.0% |
| % Total calculated<br>digestible energy from<br>protein | 17.0% | 21.0% |
| **Ingredients** |  |  |
| Casein (Acid) | 195 g/Kg | 195 g/Kg |
| Sucrose | 341 g/Kg | 341 g/Kg |
| Canola Oil | nil | 60 g/Kg |
| Clarified Butter (Ghee) | 210 g/Kg | nil |
| Cellulose | 50 g/Kg | 50 g/Kg |
| Wheat Starch | 154 g/Kg | 306 g/Kg |
| DL Methionine | 3.0 g/Kg | 3.0 g/Kg |
| Calcium Carbonate | 17.1 g/Kg | 17.1 g/Kg |
| Sodium Chloride | 2.6 g/Kg | 2.6 g/Kg |
| AIN93 Trace Minerals | 1.4 g/Kg | 1.4 g/Kg |
| Potassium Citrate | 2.6 g/Kg | 2.5 g/Kg |
| Potassium Dihydrogen<br>Phosphate | 6.9 g/Kg | 6.9 g/Kg |
| Potassium Sulphate | 1.6 g/Kg | 1.6 g/Kg |
| Choline Chloride (75%) | 2.5 g/Kg | 2.5 g/Kg |
| SF00-219 Vitamins | 10 g/Kg | 10 g/Kg |
| Cholesterol | 1.5 g/Kg | 1.5 g/Kg |
| Oxicap E2 | 0.04 g/Kg | 0.04 g/Kg |

**Table S4:**
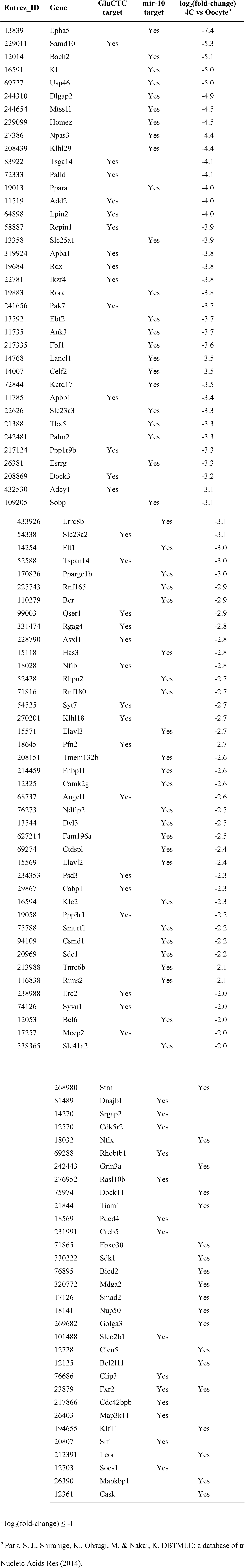
Genes targeted by tRF-GluCTC or miR-10 downregulated^a^ in the murine MZT.

